# The conserved transcriptional regulator CdnL is required for metabolic homeostasis and morphogenesis in *Caulobacter*

**DOI:** 10.1101/557637

**Authors:** Selamawit Abi Woldemeskel, Laura Alvarez, Allison K. Daitch, Rilee Zeinert, Anant Bhargava, Diego Gonzalez, Justine Collier, Peter Chien, Felipe Cava, Erin D. Goley

## Abstract

Bacterial growth and division require regulated synthesis of the macromolecules used to expand and replicate components of the cell. Transcription of housekeeping genes required for metabolic homeostasis and cell proliferation is guided by the sigma factor σ^70^. The conserved CarD-like transcriptional regulator, CdnL, associates with promoter regions where σ^70^ localizes and stabilizes the open promoter complex. However, the contributions of CdnL to metabolic homeostasis and bacterial physiology are not well understood. Here, we show that *Caulobacter crescentus* cells lacking CdnL have severe morphological and growth defects. Specifically, *ΔcdnL* cells grow slowly in both rich and defined media, and are wider, more curved, and have shorter stalks than WT cells. *ΔcdnL* cells also have aberrant localization of MreB and CtpS, cytoskeletal proteins required for maintaining proper cell width and curvature and whose function is dependent on metabolites ATP and CTP respectively. These defects arise from transcriptional downregulation of most major classes of biosynthetic genes, leading to significant decreases in the levels of critical metabolites, including pyruvate, *α*-ketoglutarate, ATP, NAD^+^, UDP-N-acetyl-glucosamine, lipid II, and purine and pyrimidine precursors. Notably, we find that *ΔcdnL* cells are glutamate auxotrophs, and Δ*cdnL* is synthetic lethal with other genetic perturbations that limit glutamate synthesis and lipid II production. Our findings implicate CdnL as a global regulator of genes required for metabolic homeostasis that impacts morphogenesis through availability of lipid II and through metabolite-mediated changes in localization of cytoskeletal regulators of cell shape.

## Main

In order to replicate, bacterial cells must synthesize enough macromolecules to double in size using nutrients available in their environment. These macromolecules are used to build structural components of the cell such as the membrane and cell wall, to synthesize the multitude of enzymes required for essential biochemical processes, and to duplicate genetic material. Transcriptional control of housekeeping genes encoding factors that carry out these essential functions contributes to maintaining metabolic homeostasis to support growth and development. Transcription initiation of housekeeping genes is achieved by housekeeping sigma factors that direct the core RNA polymerase (RNAP) to their promoter regions^1^, but may be co-regulated by other factors, including the CarD-like transcriptional regulator CdnL. CdnL is broadly conserved in bacteria and is best-characterized in *Myxococcus xanthus* and in *Mycobacteria*^2–6^. It localizes to promoter regions where the housekeeping sigma factor resides, directly binds to the RNAP *β* subunit, stabilizes the open promoter complex, and is required for transcription from rRNA promoters^2, 3, 5, 7^. *Mycobacterium tuberculosis* CarD (the CdnL homolog) is essential for growth in culture, persistence in mice, and survival during genotoxic stress and starvation^2, 4^. CdnL is also essential in *M. xanthus*, where its depletion causes cell filamentation^6^. This last observation implies that CdnL-dependent changes in gene expression are required to support proper morphogenesis, at least in *M. xanthus*.

Bacterial morphology is maintained by the peptidoglycan (PG) cell wall, which must be expanded and remodeled in a regulated manner to allow growth and division. It lies just outside the inner membrane and is a continuous, covalently-linked structure made of glycan strands crosslinked together by peptide side chains^8^. In addition to giving the cell its stereotyped shape, the cell wall provides physical integrity and prevents cell lysis due to turgor pressure. Different shape features (e.g. cell length, width, or curvature) are specified by spatially and temporally regulated synthesis and remodeling of the PG that is largely orchestrated by cytoskeletal proteins^8, 9^. Though the details of PG synthesis and remodeling and its regulation by cytoskeletal proteins has been the subject of intense study for several decades, how PG metabolism is regulated in coordination with nutrient availability and metabolic status of the cell is incompletely understood.

Here, we investigate how CdnL affects metabolism, growth, and morphogenesis in the α-proteobacterium *Caulobacter crescentus* (hereafter *Caulobacter*). *Caulobacter* undergoes stereotyped morphological transitions as it progresses through the cell cycle and during adaptation to a variety of stresses, making it an ideal organism to study bacterial morphogenesis^10^. Although it is known that *Caulobacter* CdnL is not required for viability, the consequences of its loss on *Caulobacter* physiology have not been previously characterized^11^. Here, we demonstrate that CdnL regulates transcription of biosynthetic genes required for metabolic homeostasis that, in turn, supports proper growth, PG metabolism, and morphology in *Caulobacter*.

## Results

### Δ*cdnL* cells have growth and morphology defects

We became interested in *Caulobacter* CdnL through a screen for spontaneous suppressors of a dominant lethal mutant of the cell division protein FtsZ (called FtsZΔCTL) that causes lethal defects in PG metabolism^12^. Specifically, one of the FtsZΔCTL suppressors we identified carried a point mutation in *cdnL* (*CCNA_00690*) encoding an I42N missense mutation that disrupted CdnL protein stability (Supplementary figure 1a). To examine the role of *cdnL* in *Caulobacter* growth and morphogenesis, we deleted it in a clean genetic background and compared growth and morphology to wild type (WT) cells. We found that Δ*cdnL* cells grew more slowly than WT (Figure 1a) and had pleiotropic morphological defects in complex PYE medium (Figure 1b, c). Quantitative analysis of shape differences between WT and Δ*cdnL* cells grown in PYE using CellTool^13^ revealed significant differences in two shape modes which approximately correspond to cell curvature (shape mode 2) and cell width (shape mode 3) (Figure 1c). Specifically, we observed that in PYE, *ΔcdnL* cells were wider and more curved than WT cells (Figure 1b, c, d). In these shape modes, Δ*cdnL* cells were also more variable in their range of values than WT, indicating less precise maintenance of shape in the mutant (Figure 1c). Additionally, we observed phase-light “ghost” cells in Δ*cdnL* samples indicative of cell lysis and suggesting a lack of integrity in the cell envelope (Figure 1b, arrows).

**Figure 1:**
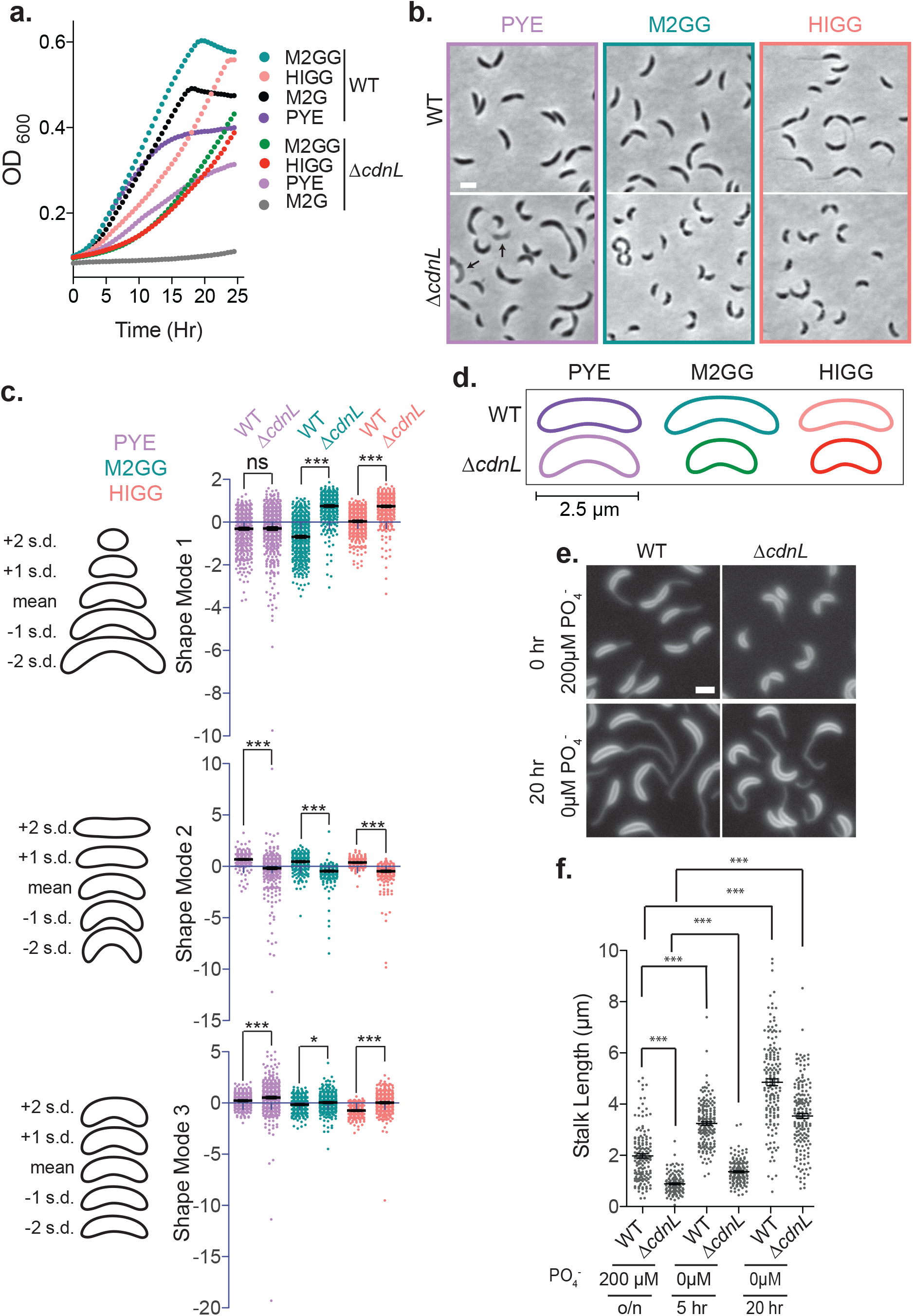
Cells lacking CdnL have pleiotropic morphological and growth defects. **a**. Growth of WT (EG865) and *ΔcdnL* (EG1447) cells in the indicated media. **b.** Phase contrast images of WT and *ΔcdnL* cells grown in the indicated media. Arrows indicate ghost cells. Bar = 2 µm. **c.** PCA of cell shape for WT and Δ*cdnL* shown in **b.** Scatter plots of shape modes 1, 2 and 3 approximately correspond to length, curvature and width. Contours on the left of scatter plots indicate mean cell shape and standard deviations (s.d) from the mean. **d.** Mean shapes of WT and Δ*cdnL* cells grown in the indicated media. **e.** Membrane staining of WT (EG865) and Δ*cdnL* (EG1447) cells with FM4-64 after growth in HIGG media with and without phosphate. **f.** Stalk lengths measured using ImageJ’s^44^ line tool. Mean and SEM included for plots in **c.** and **f.** Statistical analysis in **c.** and **f.** performed using one-way ANOVA with Bonferroni’s Multiple Comparison Test, n = 500 for each in **c.** n = 150, 127, 168, 155, 162, 172 from left to right in **e.** *** = p < 0.001, * = p < 0.05, ns = not significant.

To further characterize the Δ*cdnL* growth and morphology phenotypes, we grew WT and Δ*cdnL* cells in the defined media M2G or HIGG. Surprisingly, we found that M2G was unable to support growth of cells lacking CdnL, but that Δ*cdnL* cells grew in HIGG, albeit more slowly than WT (Figure 1a). A comparison of the components of M2G and HIGG media suggested that the missing nutrient in M2G that is required for growth of Δ*cdnL* cells might be glutamate. Consistent with this, addition of sodium glutamate to M2G (called M2GG with sodium glutamate added) supported growth of Δ*cdnL* cells to a similar rate as in HIGG. These data indicate that Δ*cdnL* cells are unable to synthesize sufficient glutamate to support growth and are now glutamate auxotrophs (Figure 1a). No other amino acids are included in either M2G or HIGG, suggesting a specific requirement for exogenous glutamate.

Similar to our observations with PYE, we found that Δ*cdnL* cells grow more slowly than WT in M2GG or HIGG and also had aberrant shape (Figure 1a, b). Δ*cdnL* cells were again hypercurved and wider than WT cells, but unlike in PYE, Δ*cdnL* cells were also consistently shorter than WT in each defined medium (Figure 1b, c, d). The polar stalk is a prominent morphological feature of *Caulobacter* cells and, like other features of cell shape, stalk biogenesis and elongation requires PG synthesis^14^. Since we observed stereotyped shape changes in cells lacking CdnL, we also compared stalk morphogenesis between WT and Δ*cdnL* cells. We found that in phosphate replete HIGG media (containing 200 µM PO_4_^-^) Δ*cdnL* cells had short stalks compared to WT cells (Figure 1e, f), suggesting that CdnL impacts all aspects of morphogenesis in *Caulobacter*. Stalks elongate in response to phosphate starvation in *Caulobacter* through a pathway that is distinct from developmentally regulated stalk morphogenesis. Interestingly, Δ*cdnL* cells significantly elongated their stalks during phosphate starvation (Figure 1 e, f) suggesting that Δ*cdnL* cells can still alter their morphology in response to phosphate starvation.

### MreB is hyperfocused at midcell in Δ*cdnL* cells

In bacteria, cytoskeletal proteins spatially regulate cell wall synthesis required for shape determination and maintenance^10^. To investigate if the shape defects that we observe in Δ*cdnL* arise due to changes in levels of cytoskeletal proteins, we probed for changes in levels of MreB, FtsZ, crescentin, and CTP synthase and found no differences in protein or transcript levels between Δ*cdnL* and WT cells (Supplementary table 1 and data not shown).

Since Δ*cdnL* cells had significant differences in width and curvature, we focused on the cytoskeletal proteins that determine these shape features. In bacteria, the actin homolog and ATPase MreB is required for rod shape, width maintenance, and cell body elongation^15, 16^. Additionally, MreB influences curvature of *Caulobacter* cells through its effects on the intermediate filament-like protein, crescentin^17^. To determine if the localization of MreB is altered in Δ*cdnL*, we expressed *venus-mreB* from the *xylX* promoter and visualized MreB localization in WT or Δ*cdnL* cells grown in PYE, M2G, and HIGG media (Figure 2 a,b,c). In WT cells grown in PYE, MreB has a patchy localization along the length of the cell in newborn swarmer (G1) cells, shifts to midcell in stalked (S) and early predivisional cells, and assumes a patchy localization in late predivisional cells^15^. Interestingly, we found that the localization of MreB at midcell is more tightly focused in Δ*cdnL* cells compared to WT in all three media tested (Figure 2a,b,c). Moreover, we found that most WT cells localize MreB along the length of the cell when grown in HIGG media (Figure 2c). However, the localization of MreB in HIGG shifts to mostly at midcell when *cdnL* is deleted. To better visualize the localization of MreB across populations and throughout the cell cycle, we created demographs. In cells that have MreB localized at midcell, we found that the localization of MreB is more tightly focused with minimal signal outside of the midcell region (Figure 2d,e,f). To quantitatively compare the differences in MreB localization between WT and Δ*cdnL*, we fit the MreB signal at midcell to an eight term Fourier series model and calculated the full width at half max (FWHM). We found that the FWHM at midcell is smaller in Δ*cdnL* cells compared to WT, consistent with what we inferred from the images and demographs (Figure 2g,h,i).

**Figure 2:**
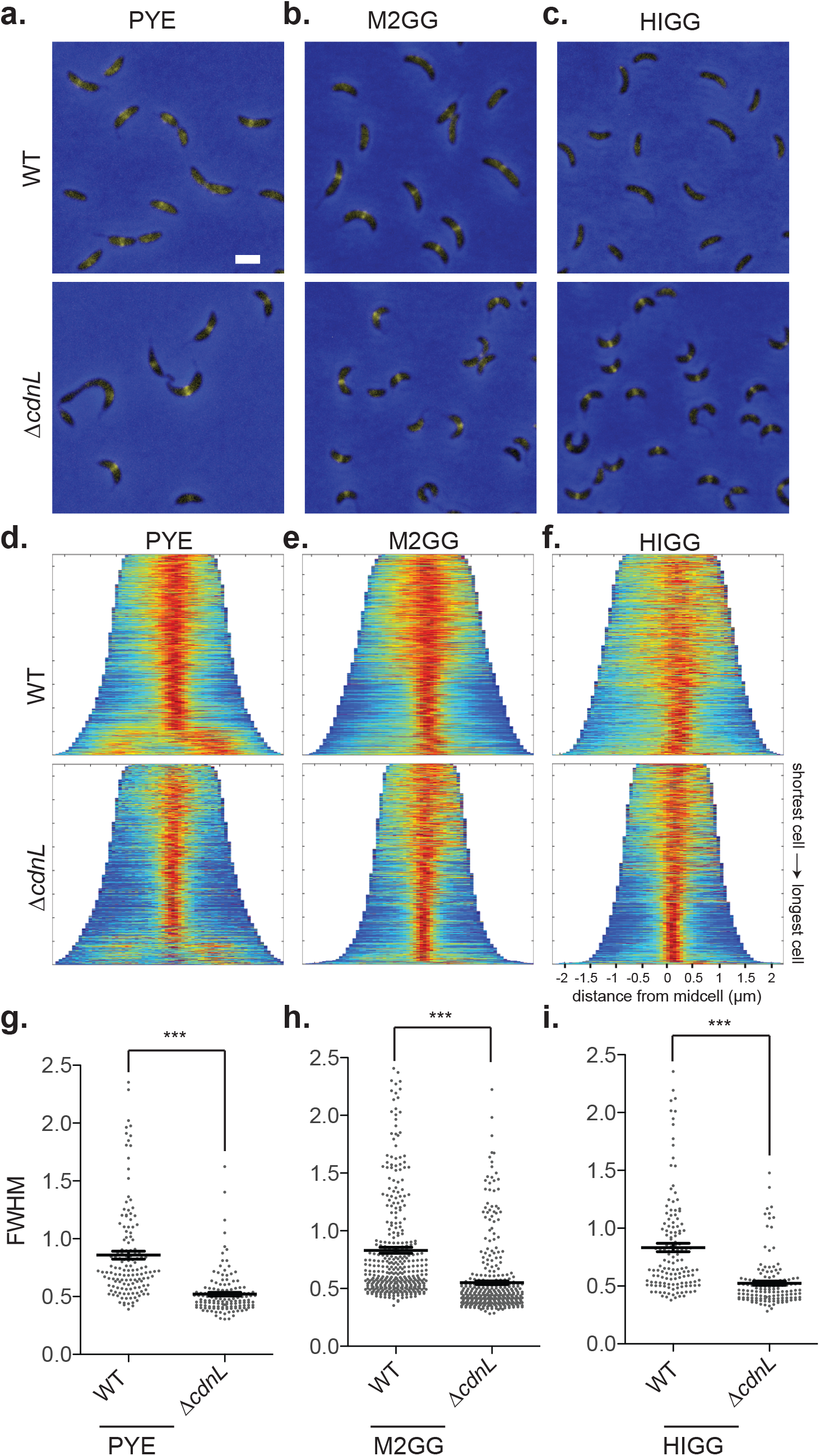
MreB localization is aberrant in Δ*cdnL* cells. **a, b & c.** Merged phase contrast (blue) and fluorescence (yellow) images of WT (EG1795) and Δ*cdnL* cells (EG2465) expressing *venus-mreB* for 2 hrs from the *xylX* promoter in the indicated media. Bar = 2 µm. **d, e & f.** Demographs of images shown in **a, b &c.** n = 850 cells for each. **g, h & i.** Scatter plots of full width at half maximum (FWHM) values of midcell-localized MreB. Cell length range used for FWHM calculation of midcell-localized MreB was determined from demographs shown in **d-f**. For PYE, WT and Δ*cdnL*, cells 2 – 3 µm long have MreB localized at midcell and were selected for FWHM calculations. For M2GG, WT and Δ*cdnL* cells between lengths of 3 – 3.75 µm and 2 – 3 µm, respectively, were selected for FWHM calculations. For HIGG, WT and Δ*cdnL* cells between lengths of 3 – 3.5 µm and 2 – 3 µm, respectively, were selected for FWHM calculations. Statistical analysis performed using unpaired t test. *** = p < 0.001. n = 140 each for PYE and HIGG and n = 383 each for M2GG.

To further characterize the link between MreB localization and loss of *cdnL,* we deleted *cdnL* in backgrounds that have mutations near the ATP binding pocket of MreB. These mutations cause MreB to localize at the poles, along the length of the cell, or at midcell throughout the cell cycle^17^. These aberrant localizations of MreB cause changes in cell shape but do not affect growth rate (Supplementary figure 2a,b,c). Deletion of *cdnL* in each of these backgrounds caused severe morphological and growth defects (Supplementary figure 2a,b). The severity of growth and shape defects was worse in mutant backgrounds that do not localize MreB to midcell (Supplementary figure 2). Interestingly, we found that MreB V324A, which has a patchy localization in a WT background, becomes enriched at midcell when *cdnL* is deleted and grows similarly to mutants that localize MreB to midcell (Supplementary figure 2b,c,d,e). These data suggest that localization of MreB at midcell is important for fitness in Δ*cdnL* cells. Since MreB is essential for specification and maintenance of width and curvature, the mislocalization of MreB may contribute to the width and curvature defects that we observe in Δ*cdnL*.

### Hypercurvature of Δ*cdnL* cells is dependent on crescentin and may involve CTP synthase

In *Caulobacter*, crescentin drives the patterning of PG synthesis required for cell curvature^9, 18^. To determine if hypercurvature of Δ*cdnL* cells is dependent on crescentin, we deleted *cdnL* in a Δ*creS* background. We observed that Δ*cdnL*Δ*creS* cells are straight, indicating that hypercurvature is dependent on crescentin (Figure 3a,b). In addition to crescentin, curvature is influenced by the metabolic enzyme CTP synthase (CtpS), which forms filaments near crescentin when CTP is abundant and negatively regulates curvature in a crescentin-dependent manner^19, 20^. Since we observed that Δ*cdnL* cells are hypercurved, we hypothesized that the hypercurvature of Δ*cdnL* may arise due to fewer CtpS filaments. Thus, we expressed *mcherry-ctpS* from the xylose-inducible promoter in WT or Δ*cdnL* cells and visualized localization of CtpS filaments and found that Δ*cdnL* cells form fainter CtpS filaments compared to WT (Supplementary figure 3). Additionally, demograph analysis of mCherry-CtpS localization indicated that mCherry-CtpS forms less robust filaments and has more dispersed signal in Δ*cdnL* cells grown in PYE or M2GG compared to WT (Figure 3c, Supplementary figure 3). Thus, our data suggest that formation of fewer CtpS filaments in Δ*cdnL* cells may lead to the hypercurvature phenotype that we observe.

**Figure 3:**
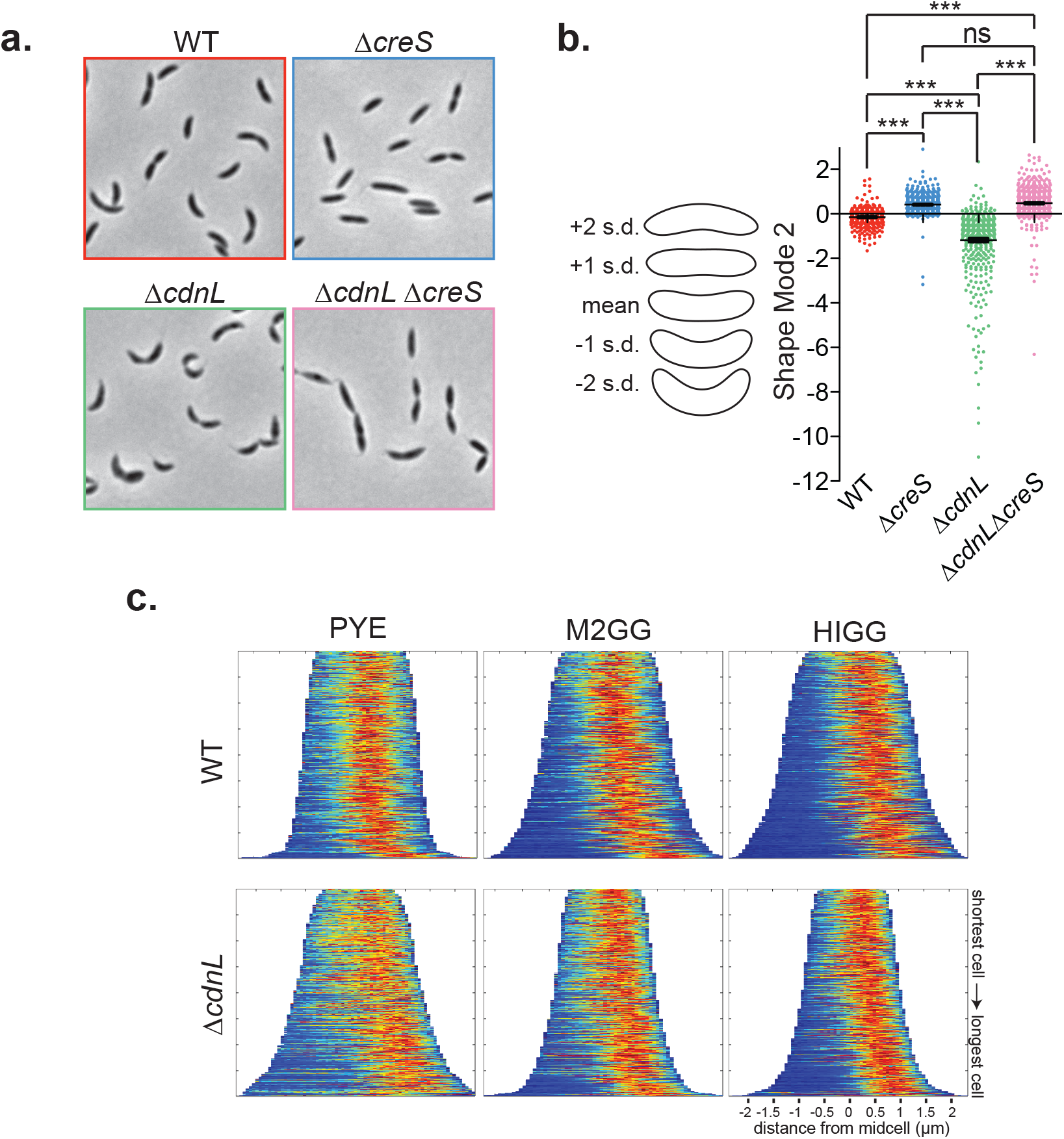
Hypercurvature of Δ*cdnL* cells is dependent on crescentin and may involve CTP synthase. **a.** Phase contrast images of WT (EG865), Δ*creS* (EG1903), *ΔcdnL::gentR* (EG1898) and Δ*creS* Δ*cdnL::gentR* (EG1912). **b.** Scatter plot depicting shape mode 2 (∼curvature) from PCA of cell shape for cells shown in **a**. Mean and SEM values are indicated. n = 447, 615, 734, 574 from left to right. Contours on the left of scatter plots indicate mean cell shape and standard deviations (s.d) from the mean. Statistical analysis performed using one-way ANOVA with Bonferroni’s Multiple Comparison Test. *** = p < 0.001, ns = not significant. **c.** Demographs of images from Supplementary Figure 3 of WT (EG1902) and Δ*cdnL* (EG1958) cells expressing *mCherry – ctpS* from the *xylX* locus grown in indicated media. n = 400 cells for each.

### Genes involved in biosynthetic processes are downregulated in Δ*cdnL*

Having established that the transcriptional regulator CdnL plays a role in growth and morphogenesis and influences the localization of morphogenetic proteins, we next sought to understand how it exerts these effects. To identify genes that are misregulated in Δ*cdnL* that may be responsible for the growth and morphology defects that we observe, we extracted mRNA from WT and Δ*cdnL* cells and performed RNA-Seq. The results from this analysis showed that approximately 30% of the transcriptome is misregulated in cells lacking CdnL (Supplementary table 1). A closer look at genes that are downregulated in the Δ*cdnL* clone that we used for RNA-seq analysis suggested that a 50 kbp region comprising 51 genes between CCNA_02773 and CCNA_02826 was highly downregulated in Δ*cdnL*. This region is flanked by identical sequences that encode transposases CCNA_02772 and CCNA_02828, suggesting that this region may be deleted in the Δ*cdnL* clone (EG1415) used in our RNA-seq analysis. Indeed, deletion of this region was confirmed by PCR. However, we found no differences in growth or morphology between the Δ*cdnL* clone with the 50 kbp deletion (EG1415, Δ50kbp) and Δ*cdnL* clones containing this region (e.g. EG1447), indicating that this region does not contribute significantly to the growth and shape defects that we observe (Supplementary figure 1b, c). Previously, this region of the chromosome was shown to be readily lost in a background lacking the methyltransferase *ccrM*^21^. Similar to our observations with Δ*cdnL*, loss of this region in Δ*ccrM* did not yield any specific growth advantages. To ensure that loss of this region did not impact our analysis of the transcriptional consequences specific to deleting *cdnL*, we analyzed the transcriptome of an independent Δ*cdnL* clone that has the 50 kbp region intact (JC784) by microarray analysis (Supplementary table 1). We found significant overlap between our RNA-Seq dataset using EG1415 and microarray dataset using JC784 (Supplementary figure 1d, e, f, g). We generally used the intersection of our two datasets (RNA-Seq of EG1415 and microarray of JC784) as a conservative and rigorous representation of the consequences of deleting *cdnL* on the *Caulobacter* transcriptome.

To obtain an overview of how loss of CdnL affects the transcriptome we used DAVID^22^ functional annotation analysis to functionally categorize genes that had at least a two-fold change in transcript abundance in Δ*cdnL* in both the RNA-Seq and microarray datasets (Supplementary table 2). Of the genes that are downregulated in Δ*cdnL*, we found an overrepresentation of genes involved in biosynthetic and bioenergetic pathways including amino acid, nucleotide, fatty acid, lipid, cell wall, cell membrane, and central carbon metabolism. Genes that are over 2-fold upregulated in Δ*cdnL* clustered into categories involved in transcription, transport, and motility.

### Δ*cdnL* cells have reduced levels of central carbon and TCA metabolites required for synthesizing macromolecules critical for growth

Since our transcriptome analysis revealed downregulation of pathways involved in macromolecular biosynthesis, we hypothesized that Δ*cdnL* cells may have limited amounts of substrates available for proliferative processes. To identify how the metabolome changes when *cdnL* is deleted, we extracted metabolites from WT and Δ*cdnL* cells grown in PYE, M2GG, M2G (grown in M2GG, washed and grown in M2G for 12 hours) or HIGG media or CdnL*-* depleted cells grown in PYE and performed LC/MS analysis of polar metabolites. Consistent with what we inferred from our transcriptomic data, we found glycolytic and tricarboxylic acid (TCA) intermediates, amino acids, nucleotides, and their derivatives to be significantly altered, suggesting that central carbon metabolism and carbohydrate utilization is reduced in cells lacking CdnL (Figure 4, Supplementary table 3 & Supplementary figure 4). Specifically, we found biosynthetic precursors and cofactors such as pyruvate, NAD^+^, ATP, and UDP-N-acetyl glucosamine are significantly reduced in Δ*cdnL* cells while D-xylose, uric acid, the TCA intermediates fumarate and malate, the pyrimidine and amino acid precursor dihydroorotate, the leucine synthesis intermediate 2-isopropylmalic acid, and the storage molecule 3-hydroxybutyric acid are significantly higher in Δ*cdnL* across all conditions compared to WT (Figure 4, Supplementary table 3 & Supplementary figure 4). The presence of higher uric acid across all media suggests that Δ*cdnL* cells are preferentially catabolizing amino acids rather than carbohydrates as a carbon source.

**Figure 4:**
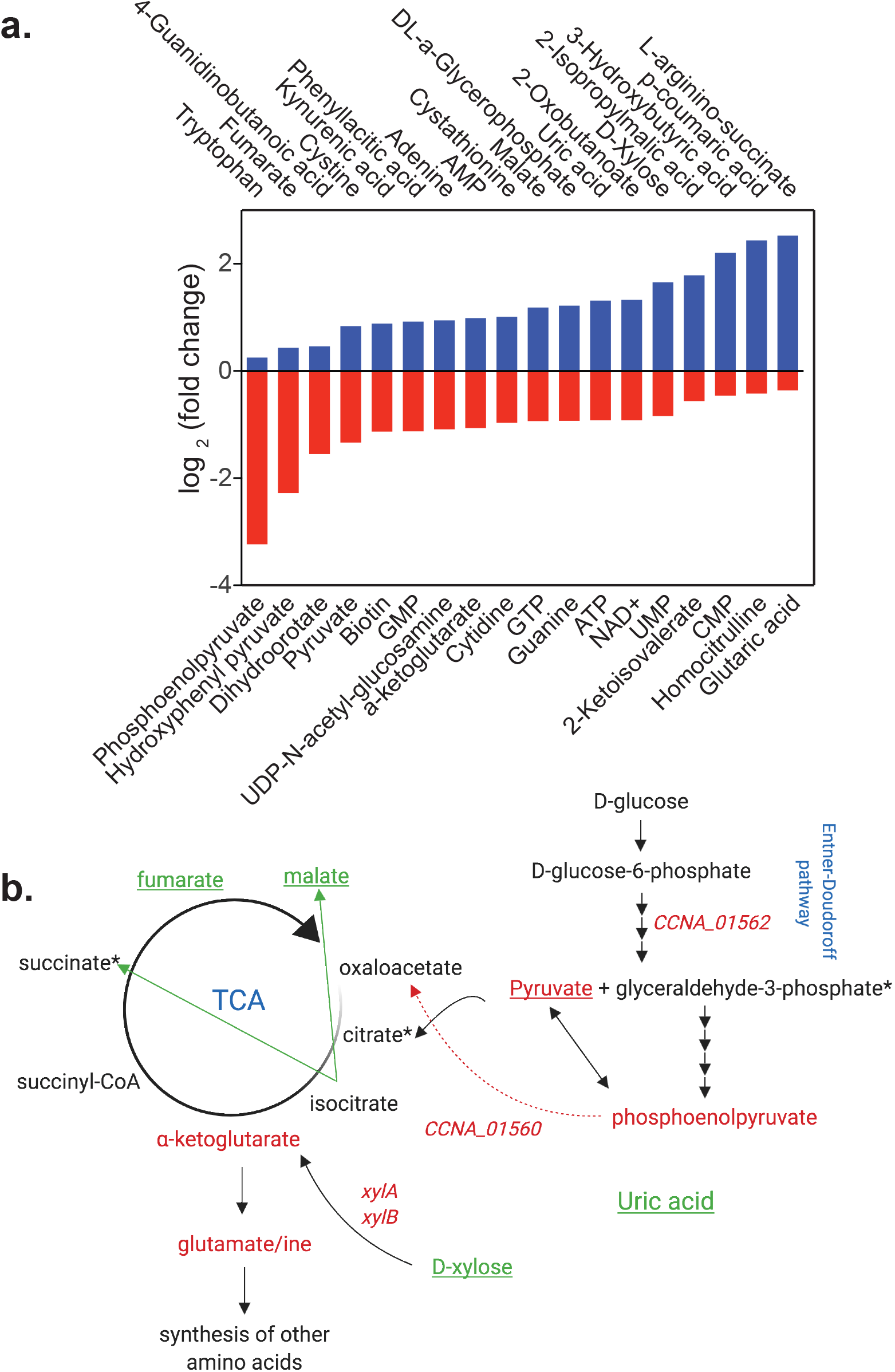
CdnL is required for metabolic homeostasis in *Caulobacter*. **a**. Graph showing significantly altered metabolites (p < 0.05) in Δ*cdnL* compared to WT grown in PYE from Supplementary table 3. **b**. Summary figure depicting changes in glucose metabolism and TCA cycle in Δ*cdnL* cells as compared to WT. Genes and metabolites shown in red are downregulated and in green are upregulated. Underlined metabolites are changed across all media types, whereas those in black are unchanged. * indicates metabolites not detected. n = 3 for each strain in each condition.

Since we performed our transcriptomics analysis in PYE, we compared our RNA-Seq data to our PYE metabolome dataset. In *Caulobacter*, glucose is metabolized to yield substrates that feed into the TCA cycle via the Entner-Doudoroff pathway^23, 24^. Since intermediates in and products of this glycolytic pathway are reduced in Δ*cdnL*, we looked at our transcriptomic data to see if the levels of transcripts encoding any of the enzymes in this pathway are changed. We found that *CCNA_01562* and *CCNA_01560* are significantly downregulated suggesting that disruption in flux through the Entner-Doudoroff pathway may, indeed, lead to low levels of pyruvate and phosphoenolpyruvate (PEP) (Figure 4b, Supplementary table 1). WT *Caulobacter* can synthesize all amino acids *de novo* using TCA cycle intermediates^23^. During growth in media without amino acids, *Caulobacter* cells can convert PEP to oxaloacetate to replenish TCA intermediates^23^. We found that *CCNA_01560*, which converts PEP to oxaloacetate, is also highly downregulated in Δ*cdnL*. Thus, our observations of low *α*-ketoglutarate levels and glutamate auxotrophy may arise due to reduced flux of pyruvate and PEP into the TCA cycle (Figure 4). Interestingly, D-xylose levels were elevated in Δ*cdnL* cells in all media tested, although xylose was not provided exogenously. Conversely, genes required to metabolize xylose are downregulated in Δ*cdnL*. Previously, xylose accumulation without an ability to metabolize xylose has been shown to upregulate isocitrate lyase (*CCNA_01841*) and initiate the glyoxylate bypass which promotes conversion of isocitrate to succinate, bypassing key TCA cycle intermediates such as *α*-ketoglutarate.^23, 25^ Additionally, upregulation of malate synthase (*CCNA_01843*) combines glyoxylate produced by isocitrate lyase to acetyl-coenzyme A to produce malate, allowing a modified TCA cycle to continue.^23^ Consistently, we find that *CCNA_01841* and *CCNA_01843* are over four-fold and two-fold upregulated in Δ*cdnL*, respectively (Supplementary table 1), suggesting low levels of *α*-ketoglutarate may arise due to activation of the glyoxyate bypass in addition to low levels of pyruvate and PEP. Since *α*-ketoglutarate is used to synthesize glutamate (Figure 4b), the glutamate auxotrophy of Δ*cdnL* cells may arise due to low amounts *α*-ketoglutarate produced by the TCA cycle. Additionally, we found that *gdhA* and *gltB*, genes essential for glutamate biosynthesis from *α*-ketoglutarate or glutamine, are over 4-fold downregulated in our transcriptomic analysis (Figure 5a, Supplementary table 1). Collectively, our data indicate that changes in the transcriptome that arise when *cdnL* is deleted have detrimental consequences on metabolic pathways disrupting levels of key metabolites required for energy production and amino acid and nucleotide biosynthesis.

**Figure 5:**
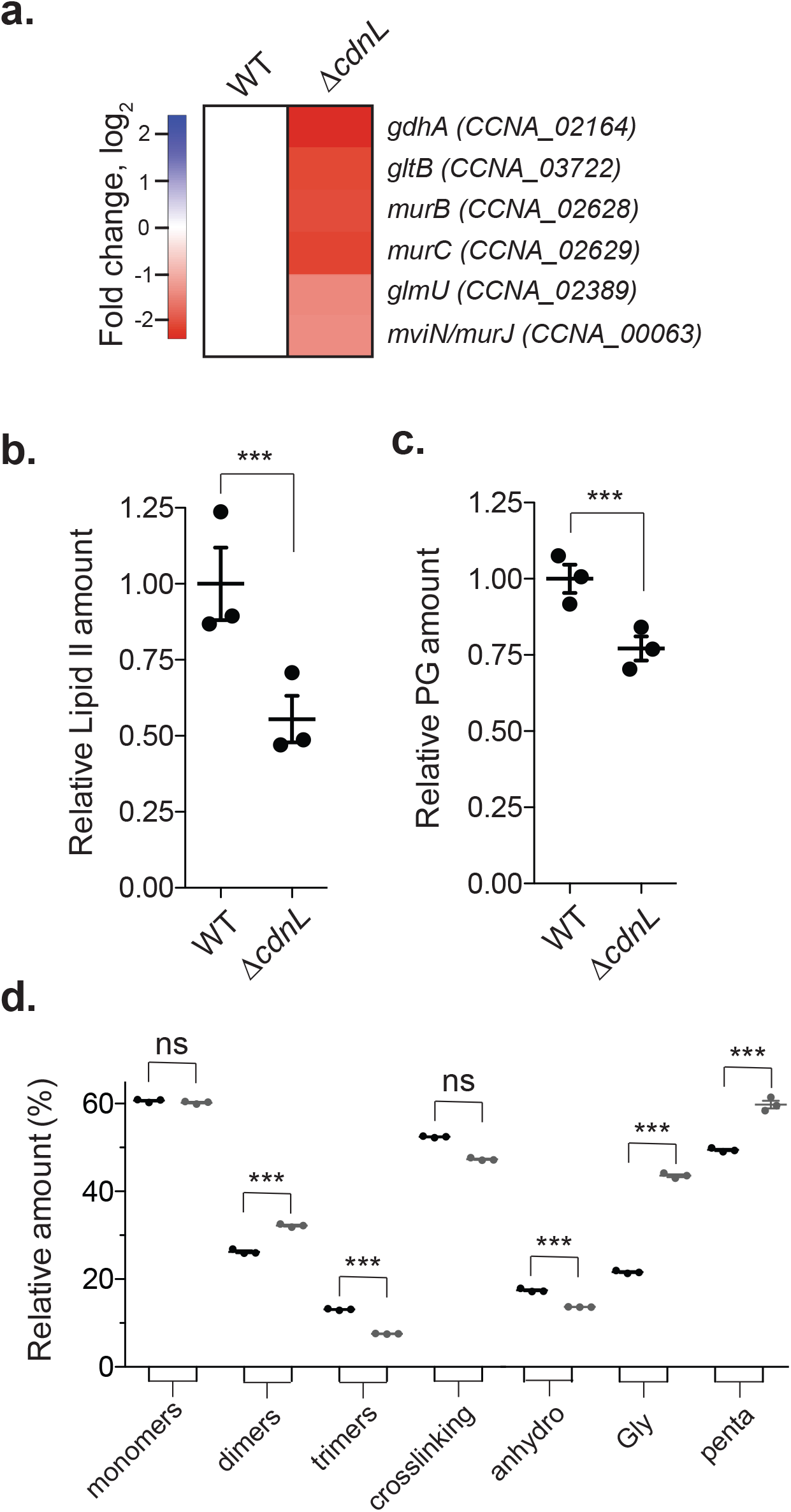
CdnL impacts cell wall precursor synthesis and cell wall metabolism. **a.** Heatmap showing changes of the indicated transcripts involved in lipid II synthesis and cell wall remodeling in WT (EG865) and Δ*cdnL* (EG1415) using RNA-Seq. **b. & c.** Relative amount of lipid II and peptidoglycan in WT (EG865) and Δ*cdnL* (EG1447) cells. **d.** Indicated muropeptide species from WT (EG865) and Δ*cdnL* (EG1447). X-axis shown in **d** are (1-6 anhydro) N-acetyl muramic acid (anhydro), Gly containing muropeptides (Gly), and pentapeptides (penta). Values in **b, c, & d** are from three independent cultures and mean and SEM are indicated. Statistical analysis performed using unpaired t test. *** = p < 0.001, ** = p < 0.01, * = p < 0.05.

### Δ*cdnL* cells have low levels of the cell wall precursor and changes in cell wall crosslinking

Our metabolomics analysis showed that UDP-N-acetylglucosamine and PEP, substrates required for synthesis of the PG precursor lipid II, are significantly lower in Δ*cdnL* compared to WT (Figure 4). Additionally, our transcriptomic analysis indicated that genes required for lipid II biosynthesis are highly downregulated (Figure 5a) leading us to postulate that PG material maybe limiting in Δ*cdnL* cells. Since PG synthesis is required for both cell shape maintenance and cell envelope integrity, changes in abundance of lipid II might underlie the shape and cell lysis phenotypes observed for Δ*cdnL* cells. We directly compared lipid II levels in WT and Δ*cdnL* cells and found that Δ*cdnL* cells have a striking 45% reduction in lipid II levels as compared to WT (Figure 5b).

To assess if CdnL-mediated changes in transcription result in alterations to PG metabolic activities in addition to lipid II synthesis, we performed muropeptide analysis on PG isolated from WT and Δ*cdnL* cells to identify changes in PG chemistry. Consistent with low levels of lipid II, Δ*cdnL* cells have 23% less PG polymer than WT (Figure 5c). Although overall PG crosslinking is unaffected, Δ*cdnL* cells had significantly more dimers and fewer trimers (higher order crosslinks) than WT (Figure 5d, Supplementary Figure 5). The mutant also shows an increased chain length, as the relative amount of anhydro muropeptides (glycan chain termini) were significantly reduced in Δ*cdnL* cells (Figure 5d, Supplementary Figure 5). Moreover, Δ*cdnL* cells had more pentapeptides and glycine-containing muropeptides compared to WT (Figure 5d, Supplementary Figure 5). Thus, the shape and envelope integrity defects observed in Δ*cdnL* cells may arise, at least in part, from a combination of low levels of lipid II and significant changes in the chemical structure of the cell wall.

### Pathways impacting PG metabolism become essential in Δ*cdnL*

To gain insight into which transcriptional and metabolic changes in Δ*cdnL* may be most relevant to fitness, we performed comparative transposon sequencing (Tn-Seq) on WT and Δ*cdnL* strains. Consistent with morphological defects and low amounts of lipid II observed for Δ*cdnL* cells, we found several normally non-essential genes involved in pathways that interact with PG synthesis could not be disrupted in the Δ*cdnL* background (Figure 6a, Supplementary table 4). Specifically, we identified genes involved in glutamate metabolism, PG metabolism, PG recycling, and capsule synthesis in which we recovered far fewer transposon insertions in the Δ*cdnL* background than in WT (Figure 6a,b, Supplementary table 4). Of the genes involved in glutamate metabolism, GdhZ and MurI become essential in Δ*cdnL*. Each of these proteins link glutamate metabolism to morphogenesis: MurI by making D-Glu for lipid II synthesis and GdhZ via its effects on the morphogenetic protein FtsZ^26, 27^.

**Figure 6:**
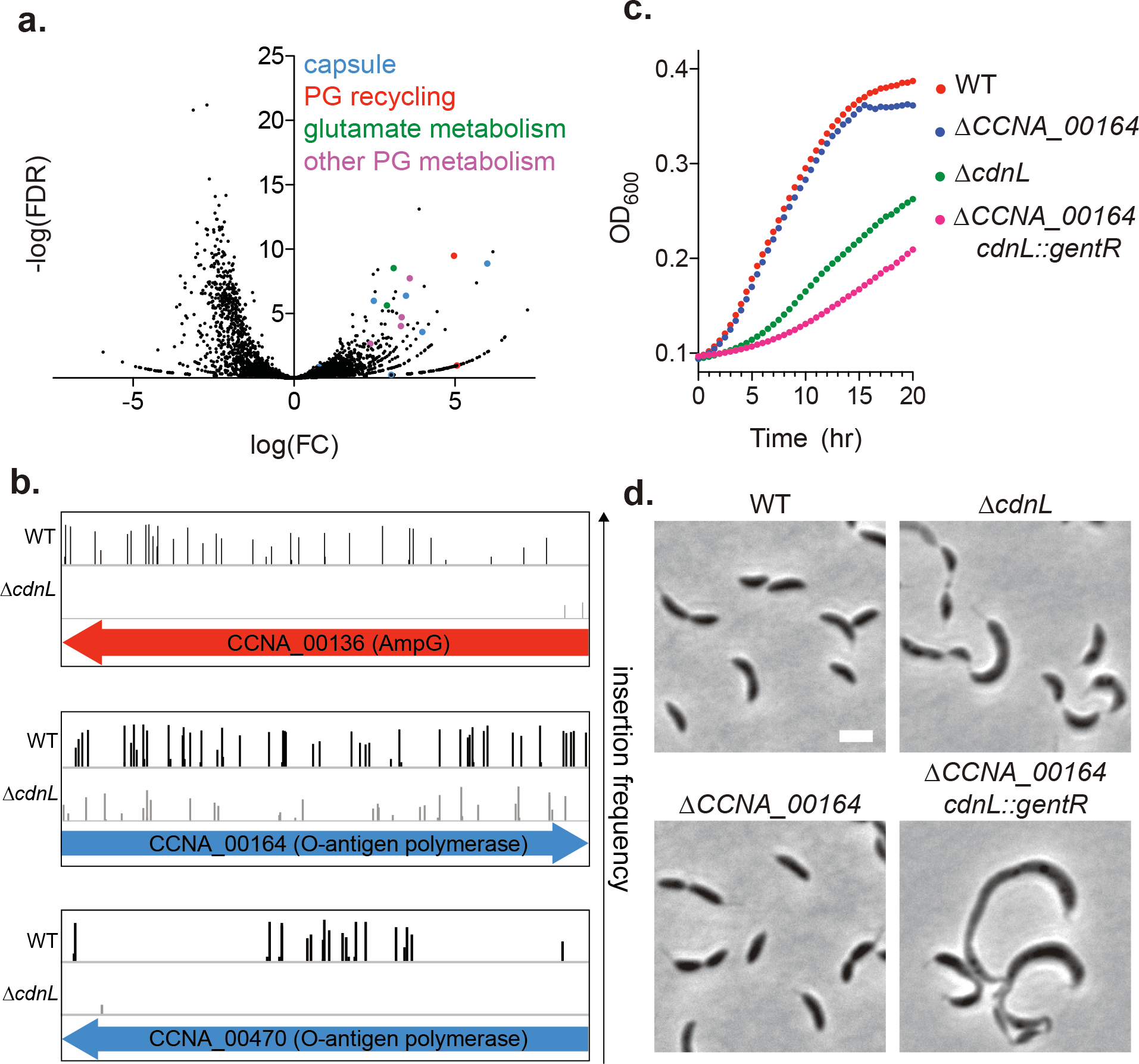
Mutations in cell wall recycling and capsule biosynthetic genes are synthetic lethal with Δ*cdnL*. **a.** Volcano plot representation of Tn-Seq analysis, WT/Δ*cdnL*. Negative log_10_ of the false discovery rate (-log(FDR)) is plotted against log_2_ of the fold change in number of unique transposon insertions in each gene in WT (EG865) vs Δ*cdnL* (EG1447). Genes that become essential in the Δ*cdnL* background and are discussed in this study are color-coded. **b.** Transposon insertion profile for selected genes from **a. c & d.** Growth curves and phase contrast images for WT (EG865), Δ*cdnL* (EG1447), *ΔCCNA_00164* (EG2604) and *cdnL::gentR ΔCCNA_00164* (EG2804). Bar = 2 µm.

Our Tn-Seq analysis also indicated that *ampG (CCNA_00136)* and *ampD* (*CCNA_02650*), genes required for PG recycling, become essential in Δ*cdnL* (Figure 6b, Supplementary table 4). AmpG is a permease that allows transport of cell wall material from the periplasm to the cytoplasm and AmpD degrades transported cell wall material to provide building blocks for lipid II synthesis^28^. We attempted to delete *cdnL* in a *ΔampG* background, but were unable to obtain clones with deletions in both *ampG* and *cdnL*, supporting the genetic interaction suggested from the Tn-Seq. Since lipid II levels are low in Δ*cdnL*, the cell apparently relies on pathways that allow for recycling of the existing cell wall material for survival.

In addition to PG metabolic genes, we found putative synthetic lethal interactions between capsule (exopolysaccharide) biosynthetic genes and *cdnL* (Figure 6a,b, Supplementary table 4). During capsule synthesis, saccharide precursors are synthesized on the lipid anchor undecaprenyl phosphate (Und-P) in the cytoplasm, flipped to the periplasmic space, and assembled into a polysaccharide that is exported to the outer surface of the cell^29^. Thus, PG synthesis and capsule synthesis each use Und-P as a lipid carrier for synthesis and export into the periplasmic space. Previous work in *Escherichia coli* and *Bacillus subtilis* has shown that disruption of pathways that use Und-P can lead to Und-P sequestration and cause severe growth and morphological defects by inhibiting lipid II synthesis^30–32^. Similarly, we propose that disrupting the capsule biosynthesis pathway in *Caulobacter* may sequester Und-P in dead-end intermediates, limiting lipid II synthesis. We attempted to delete each of the capsule genes predicted by Tn-Seq to genetically interact with *cdnL* in Δ*cdnL* cells. We were unsuccessful with the exception of *CCNA_00164*, which encodes a putative O-antigen polymerase ligase and which had the weakest predicted genetic interaction with *cdnL* of the capsule genes (Figure 6a, b and Supplementary table 4). Nevertheless, deletion of both *cdnL* and *CCNA_00164* resulted in cells with morphological defects that are more severe than Δ*cdnL* alone (Figure 6d). Moreover, these double mutant cells grew more slowly than Δ*cdnL* (Figure 6c). Deletion of *CCNA_00164* by itself had no obvious effect on growth or morphology. Our data support the model that mutations in the capsule biosynthesis pathway sequester Und-P and limit its availability for lipid II synthesis, exacerbating the existing lipid II deficiency of Δ*cdnL* cells to a lethal level. Collectively, our transcriptomic data, biochemical analyses, and genetic interaction results indicate that CdnL is required to maintain adequate availability of the PG precursor lipid II to support normal growth and morphology.

## Discussion

In this work, we unveil a new role for the conserved, global transcriptional regulator CdnL in mediating transcription of biosynthetic genes required for metabolic homeostasis in *Caulobacter*. We show that in Δ*cdnL* cells, transcription of many classes of biosynthetic genes is downregulated, with physiologically critical defects in glutamate and lipid II metabolism, in particular. These changes in metabolic pools affect growth and morphology directly by limiting macromolecules required for cell growth and envelope expansion, and indirectly by influencing morphogenetic proteins. Downregulation of genes involved in biosynthetic and bioenergetics pathways in Δ*cdnL* cells, an altered metabolic landscape in Δ*cdnL*, and the reliance of Δ*cdnL* cells on exogenous glutamate, which is a central node in multiple biosynthetic pathways, establish CdnL as a global regulator of genes required for central metabolism and macromolecular synthesis^33, 34^.

Previously, CdnL has been shown to directly bind to RNAP and stabilize the open promoter complex required for transcription of rRNA promoters in *Mycobacteria* and *M. xanthus*^3, 5^. Similarly, *Caulobacter* CdnL has been shown to bind RNAP and regulate transcription from an rRNA promoter^11^. Here, we demonstrate that loss of CdnL has global effects on various aspects of bacterial metabolism. Interestingly, however, transcriptomic changes that occur during loss of CdnL in *Caulobacter* are, in many aspects, the opposite of what is observed in *Mycobacterium smegmatis*. CarD depletion in *M. smegmatis* leads to upregulation of genes encoding the translation machinery and enzymes in metabolic pathways such as amino acid biosynthesis and to downregulation of genes involved in degradation of amino acids, fatty acid metabolism and membrane proteins^2^. Although loss of CdnL generally has opposite effects on expression of metabolic genes in these two divergent bacteria, CdnL-mediated effects on the transcriptome impact proliferative and metabolic processes and are thereby essential for proper growth in *Caulobacter* and viability in *Mycobacteria*.

In *Caulobacter*, loss of CdnL causes aberrant morphogenesis, at least in part, by limiting production of lipid II. The effects of *cdnL* deletion are in some ways reminiscent of loss of the master regulator and RNA chaperone Hfq in *Caulobacter*^35^. Deletion of *hfq* alters the levels of several metabolites, including glutamate and components of the TCA cycle. Importantly, this includes accumulation of *α*-ketoglutarate which, in turn, causes reduced synthesis of lipid II and pleiotropic cell shape defects^35^. Though both Hfq and CdnL are required to maintain sufficiently high levels of lipid II to support growth and stereotyped cell shape, CdnL has a distinct shape phenotype, likely mediated by additional impacts on MreB and CtpS.

In recent years, a growing number of factors have been described that link the metabolic status of the cell to morphology through cytoskeletal proteins ^8, 34, 36^. The best-studied examples of this employ moonlighting functions of metabolic enzymes to couple metabolism to morphology. For example, the glucosyltransferases UgtP and OpgH in *B. subtilis* and *E. coli,* respectively, alter FtsZ dynamics and affect cell size in a UDP-glucose dependent manner^37, 38^. Similarly, in *Caulobacter* GdhZ and KidO alter FtsZ assembly in a glutamate-dependent manner, coupling metabolism to cell division^26, 39^. In the case of CdnL, its impact on the localization of the cytoskeletal proteins MreB and CtpS may be metabolite-mediated rather than protein-mediated. Polymerization of CtpS into filaments that negatively regulate cell curvature is dependent on CTP, which is predicted to be present in low amounts in Δ*cdnL* cells by both our transcriptomic and metabolomic analyses (Fig. 4a and Supplementary table 1). It is unclear how MreB localization is regulated in *Caulobacter*, though it appears to be linked to its ATP-dependent polymerization cycle^17^ and may involve cell wall precursors. Thus, we propose that the disruption of metabolic homeostasis that arises from loss of CdnL impacts morphology through metabolite-mediated effects on cytoskeletal proteins in addition to reduced levels of lipid II.

Interestingly, depletion of CdnL in *M. xanthus* results in cell filamentation, consistent with a conserved impact on morphogenesis^6^. The effects, if any, of CarD depletion in *Mycobacteria* on cell morphology have not been reported, nor have the effects of CdnL depletion on the *M. xanthus* transcriptome or metabolome. Future work in diverse organisms will reveal if the role of CdnL as a global regulator of both morphogenesis and metabolism is conserved across bacteria.

## Methods

### Bacterial strains, plasmids, growth conditions

*Caulobacter crescentus* NA1000 strains were grown in peptone yeast extract medium (PYE), M2G^40^, M2GG (M2G with 0.15% sodium glutamate), or Hutner base imidazole-buffered glucose glutamate medium (HIGG)^41^. Xylose was used at 0.3% for induction experiments. EG1415 and JC874 were used for RNA-Seq and microarray experiments, respectively. All other experiments with Δ*cdnL* were performed using EG1447 (clean, in-frame deletion of *cdnL*) or EG1898 (*cdnL* coding sequence replaced with *aacC1,* conferring gentamicin resistance) or derivatives thereof. JC874, EG1447, and EG1898 contain the 50 kb region that is deleted in EG1415. Growth curves were performed at 30 °C in a Tecan Infinite M200 Pro plate reader, 100 µL culture volume, with each strain monitored in biological triplicate at OD_600_ every 30 min with intermittent shaking. Antibiotics for growing *Caulobacter* and cloning and *E. coli* were used at concentrations in liquid (solid) media as described in Woldemeskel *et al* 2017^42^. *C. crescentus*: kanamycin 5 (25) µg/mL, gentamicin 1 (5) µg/mL. *E. coli:* ampicillin 50 (100) µg/mL, kanamycin 30 (50) µg/mL, gentamicin 15 (20) µg/mL, and spectinomycin 50 (50) µg/mL.

### Phase contrast, ensemble fluorescence microscopy and image analysis

Cells in exponential phase of growth were spotted on 1% agarose pads and imaged using a Nikon Eclipse Ti inverted microscope equipped with a Nikon Plan Fluor 100X (NA1.30) oil Ph3 objective and Photometrics CoolSNAP HQ^2^ cooled CCD camera. Chroma filter cubes ET-EYFP and ET-dsRED were used for Venus and mCherry, respectively. Images were processed using Adobe Photoshop and demographs were made using Oufti^43^. Principal component analysis to identify shape variations were performed using CellTool^13^ after phase contrast images were converted to binary masks in ImageJ^44^ and edited in Adobe Photoshop to remove overlapping cells. Full-width at half-maximum (FWHM) calculations of midcell MreB signal from Oufti^43^ output were performed using a custom MATLAB script which fit the normalized signal output from Oufti^43^ into an eighth term Fourier series model. Cells for which the fit had adjusted R square value greater than 0.85 were chosen for FWHM calculation. Cells that have midcell localized MreB were selected for FWHM determination based on cell length bins guided by demographs.

### Immunoblotting to assess CdnL levels

His_6_-SUMO-CdnL was overproduced in Rosetta (DE3) pLysS *E.coli* cells from pEG1129 by inducing for 3 h at 30 °C with 0.5 mM IPTG at OD_600_ of 0.5. Cells were harvested by centrifugation, resuspended in lysis buffer (50 mM Tris-HCl, pH 8.0, 300 mM KCl, 20 mM imidazole, and 10% glycerol), lysed with 1 mg/mL lysozyme, sonicated and centrifuged at 4 °C for 30 min at 15000 x g. Supernatant was loaded onto a HisTrap FF 1 mL column (GE Life Sciences) and eluted with lysis buffer with 300 mM imidazole, cleaved with His_6_-Ulp1 (SUMO protease) at 1:500 (protease: fusion) molar ratio overnight while dialyzing into lysis buffer. Sample was reloaded onto HisTrap FF 1mL and flow through collected to separate His_6_-SUMO and His_6_-Ulp1 from CdnL. CdnL fraction was dialyzed into PBS prior to antibody production. Cells in log phase were isolated and lysed in SDS-PAGE loading buffer by boiling for 5 min. Standard procedures were used for SDS-PAGE and transfer to nitrocellulose membrane. CdnL antisera was generated by immunizing a rabbit with CdnL purified as above (Pocono Rabbit Farm & Laboratory). Specificity was determined using cell lysates with deleted or overexpressed *cdnL*. CdnL antisera was used at 1:5000.

### RNA-Seq preparation

Cultures of three independent colonies of WT (EG865) and Δ*cdnL* (EG1415, Δ50kb) cells were grown in PYE and harvested at OD_600_ of 0.45 and total RNA was extracted using PureLink RNA Mini Kit from Thermo Fisher using the protocol provided with the kit. Briefly, cells were harvested, lysed, and homogenized using lysozyme, SDS, provided Lysis buffer, and homogenizer. Nucleic acids were extracted with ethanol and loaded onto the provided spin cartridge. DNA-free total RNA was extracted using the on-column PureLink DNAse Treatment protocol. PureLink DNAse mixture was directly added on to the spin cartridge membrane, incubated for 15 min and washed using provided wash buffers and RNA was eluted with RNAse-free water. Finally, RNA quality was quantified using a bioanalyzer and rRNA was removed using the Ribo-Zero rRNA Removal Kit (Gram-Negative Bacteria). RNA-Seq libraries were prepared using the Illumina TruSeq stranded RNA kit and sequenced on an Illumina HiSeq 2500. Data analysis was performed using the Illumina’s CASAVA 1.8.2, Bowtie2 (v 2.2.5), and Bioconductor’s DESeq package. rRNA clean up, quality control analysis, library prep, sequencing and analysis was performed by the Johns Hopkins University School of Medicine Next Generation Sequencing Center.

### Microarray preparations

Biological triplicates of WT (JC450) and Δ*cdnL* (JC784) cultures grown in PYE were harvested at OD_660_ of 0.3. RNA was extracted with Trizol, purified using Ambion PureLink RNA mini-kit according to manufacturer’s protocol. Double stranded cDNA was prepared with Roche Double Stranded cDNA synthesis kit. Microarray analysis was performed on Nimblegen custom chips following the manufacturer’s protocol for labeling, hybridization on 4-plex chips, and washing. Scanning was performed with Agilent microarray scanner at 3 µm and normalization was performed with Nimblescan implemented RMA algorithm (3 x 4-plex chips in total, all samples from the same culture conditions). The statistical analysis of significance was performed with R using the Significance of Analysis of Microarrays (SAM) method implemented in the library *siggenes.*^45^

### DAVID analysis and functional clustering

Genes with a two-fold change in transcript levels in both RNA-Seq and microarray analysis were functionally categorized using DAVID^22^ with a medium classification stringency and default parameters.

### Transposon library preparation, sequencing, and analysis

Two Tn-Seq libraries each were generated for WT (EG865) and Δ*cdnL* (EG1447). 1L PYE cultures were harvested at OD_600_ of 0.4-0.6, washed 5 times with 10% glycerol, and electroporated with the Ez-Tn5 <Kan-2> transposome (Epicentre). Cells were recovered for 90 minutes at 30 °C with shaking, and plated on PYE-Kan plates. WT libraries were grown for 5 and 6 days and Δ*cdnL* libraries were grown for 6 and 10 days. Colonies were scraped off plates, combined, resuspended to form a homogeneous solution in PYE, and flash frozen in 20% glycerol. Genomic DNA for each library was extracted from an aliquot using the DNeasy Blood and Tissue Kit (Qiagen). Libraries were then prepared for Illumina Next-Generation sequencing through sequential PCR reactions. The first PCR round used arbitrary hexamer primers with a Tn5 specific primer going outward. The second round used indexing primers with unique identifiers to filter artifacts arising from PCR duplicates. Indexed libraries were pooled and sequenced at the University of Massachusetts Amherst Genomics Core Facility on the NextSeq 550 (Illumina).

Sequencing reads were first demultiplexed by index, each library was concatenated and clipped of the molecular modifier added in the second PCR using Je^46^:

java -jar /je_1.2/je_1.2_bundle.jar clip F1=compiled.gz LEN=6

Reads were then mapped back to the *Caulobacter crescentus* NA1000 genome (NCBI Reference Sequence: NC_011916.1) using BWA^47^ and sorted using Samtools^48^:

bwa mem -t2 clipped.gz | samtools sort -@2 -> sorted.bam

Duplicates were removed using Je^46^ and indexed with Samtools^48^ using the following command: java -jar /je_1.2/je_1.2_bundle.jar markdupes I=sorted.bam O=marked.bam M=METRICS.txt

MM=0 REMOVE_DUPLICATES=TRUE

samtools index marked.bam

The 5’ insertion site of each transposon were converted into .wig files comprising of counts per position and visualized using Integrative Genomics Viewer (IGV)^49, 50^. Specific hits for each library were determined with coverage and insertion frequency using a bedfile containing all open reading frames from NC_011916.1 and the outer 20% of each removed to yield a clean and thorough insertion profile. This was determined using BEDTools^51, 52^ and the following commands:

bedtools genomecov -5 -bg marked.bam > marked.bed

bedtools map -a NA1000.txt -b marked.bed -c 4 > output.txt

Library comparisons were performed using the edgeR package in the Bioconductor suite using a quasi-likelihood F-test (glmQLFit) to determine the false discovery rate adjusted p-values reported here.

### Lipid II extraction and analysis

Precursor extraction was performed as described previously and performed in triplicates^53^. Briefly, 500 mL of WT and Δ*cdnL* were grown in PYE to OD_600_ of 0.45. Cells were harvested, resuspended in 5 mL PBS and poured into 50 mL flasks containing 20 mL CHCl_3_:Methanol (1:2). The mixture was stirred for 1 h at room temperature and centrifuged for 10 min at 4000 x g at 4 °C. The supernatant was transferred to clean 250 mL flasks containing 12 mL CHCl_3_ and 9 mL PBS, stirred for 1 h at room temperature and centrifuged for 10 min at 4000 x g at 4 °C. The interface fraction (between the top aqueous and bottom organic layers) was collected and vacuum dried. To remove lipid tail, samples were resuspended in 100 µL DMSO and 800 µL H_2_O, 100 µL ammonium acetate 100 mM pH 4.2 were added. This mixture was boiled for 30 min, dried in vacuum and resuspended in 300 µL H_2_O. Samples were analyzed by UPLC chromatography coupled to MS/MS analysis, using a Xevo G2-XS QTof system (Waters Corporation, USA). The separation method used is identical to the one used for muropeptide separation explained below. A library of compounds was used to target the identification of peptidoglycan precursors and possible intermediates, although only lipid II (lipid II-penta) was detected in these samples. Lipid II amounts were calculated based on the integration of the peaks (total area), normalized to the culture OD.

### Peptidoglycan (PG) purification and analysis

PG samples were analyzed as described previously^54, 55^. Briefly, 50 mL of WT and Δ*cdnL* cells were grown to an OD_600_ of 0.5 in PYE, harvested, and boiled in 5% SDS for 2 h. Sacculi were repeatedly washed with MilliQ water by ultracentrifugation (110,000 rpm, 10 min, 20 °C) until total removal of the detergent and finally treated with muramidase (100 µg/mL) for 16 hours at 37 °C. Muramidase digestion was stopped by boiling and coagulated proteins were removed by centrifugation (10 min, 14,000 rpm). For sample reduction, the pH of the supernatants was adjusted to pH 8.5-9.0 with sodium borate buffer and sodium borohydride was added to a final concentration of 10 mg/mL. After incubating for 30 min at room temperature, pH was adjusted to 3.5 with orthophosphoric acid.

UPLC analyses of muropeptides were performed on a Waters UPLC system (Waters Corporation, USA) equipped with an ACQUITY UPLC BEH C18 Column, 130Å, 1.7 µm, 2.1 mm × 150 mm (Waters, USA) and a dual wavelength absorbance detector. Elution of muropeptides was detected at 204 nm. Muropeptides were separated at 45 °C using a linear gradient from buffer A (formic acid 0.1% in water) to buffer B (formic acid 0.1% in acetonitrile) in an 18-minute run, with a 0.25 mL/min flow. Relative total PG amounts were calculated by comparison of the total intensities of the chromatograms (total area) from three biological replicates normalized to the same OD_600_ and extracted with the same volumes. Muropeptide identity was confirmed by MS/MS analysis, using a Xevo G2-XS QTof system (Waters Corporation, USA). Quantification of muropeptides was based on their relative abundances (relative area of the corresponding peak) normalized to their molar ratio.

### Metabolomics sample preparation and analysis

4 mL of each strain was grown to an OD_600_ of 0.3 and filtered through 0.22 µm nylon filters (Millipore GNWP04700). The filter was placed upside down in a 60 mm dish containing 1.2 mL pre-chilled quenching solution (40:40:20 Acetonitrile:methanol:H_2_O + 0.5% formic acid) and incubated at -20 °C for 15 minutes. Cells were washed off the filter by pipetting the quenching solution over the filter, transferred to chilled bead beating tubes containing 50 mg of 0.1 mm glass beads and neutralized with 100 µL 1.9M NH_4_HCO. Cells were lysed on a beat beater using a Qiagen Tissulyzer at 30Hz for 5 minutes. Samples were spun at 4 °C for 5 min at max speed and transferred to pre-chilled tubes to remove debris.

LC-MS analysis of cellular extract was conducted on Q Exactive PLUS Hybrid Quadrupole-Orbitrap mass spectrometer (Thermo Fisher Scientific) using hydrophilic interaction chromatography. The Dionex UltiMate 3000 UHPLC system (Thermo Fisher Scientific) with XBridge BEH amide column (Waters, Milford, MA) and XP VanGuard Cartridge (Waters, Milford, MA) were used for LC separation. The LC gradient, comprising solvent A (95%:5% H2O:acetonitrile with 20 mM ammonium acetate, 20 mM ammonium hydroxide, pH 9.4) and solvent B (20%:80% H2O:acetonitrile with 20 mM ammonium acetate, 20 mM ammonium hydroxide, pH 9.4), corresponded with the following solvent B percentages over time: 0 min, 100%: 3 min, 100%; 3.2 min, 90%; 6.2 min, 90%; 6.5 min, 80%; 10.5 min, 80%; 10.7 min, 70%; 13.5 min, 70%; 13.7 min, 45%; 16 min, 45%; 16.5 min, 100%. Chromatography flow rate was at 300 µL/min and injection volume 5 µL. Column temperature was maintained at 25 °C. MS scans were set to negative ion mode with a resolution of 70,000 at m/z 200, in addition to an automatic gain control target of 3 x 10^6 and scan range of 72 to 1000. Metabolite data was obtained using the MAVEN software package^56^.

## Acknowledgements

We thank members of the Goley lab for helpful discussions and input. We thank Kousik Sundararajan for identifying the initial mutant of CdnL that suppresses the FtsZΔCTL phenotype. We would like to thank Julie Theriot, Lucy Shapiro, Christine Jacobs-Wagner, Regis Hallez, and Zemer Gitai for providing materials and strains used in this project. We thank the Johns Hopkins Genetic Resources Core Facility for RNA-Seq services and analysis, Zach Pincus for guidance in using CellTool and the Metabolomics core at the Rutgers Cancer Institute of New Jersey for metabolomics analysis. This work was funded by the NIH through R01GM108640 (to EDG), R01GM111706 (to PC), T32GM08515 (training grant support of RZ), and T32GM007445 (training grant support of AKD) and Johns Hopkins Discovery Fund grant to EDG. JC is funded by Swiss National Science Foundation (SNSF) grants 31003A_140758 and 31003A_173075. Research in the Cava lab is supported by MIMS, the Knut and Alice Wallenberg Foundation (KAW), the Swedish Research Council, and the Kempe Foundation.

## Author contributions

SAW and EDG conceived the study. SAW, LA, RZ, DG, JC, PC, FC, and EDG designed the experiments. SAW performed the majority of experiments and analysis, except as noted. LA performed muropeptide analysis and lipid II quantification. SAW, AKD, RZ, and PC performed and/or analyzed Tn-Seq experiments. AB wrote custom software to analyze MreB distribution. DG performed and analyzed microarray experiments. EDG purified CdnL, generated strains, and performed CellTool analysis of shape. JC, PC, FC, and EDG supervised the research. SAW and EDG wrote the manuscript with support and editing from all authors.

