## Supplementary Figures and Legends for "The conserved transcriptional regulator CdnL is required for metabolic homeostasis and morphogenesis in *Caulobacter*"

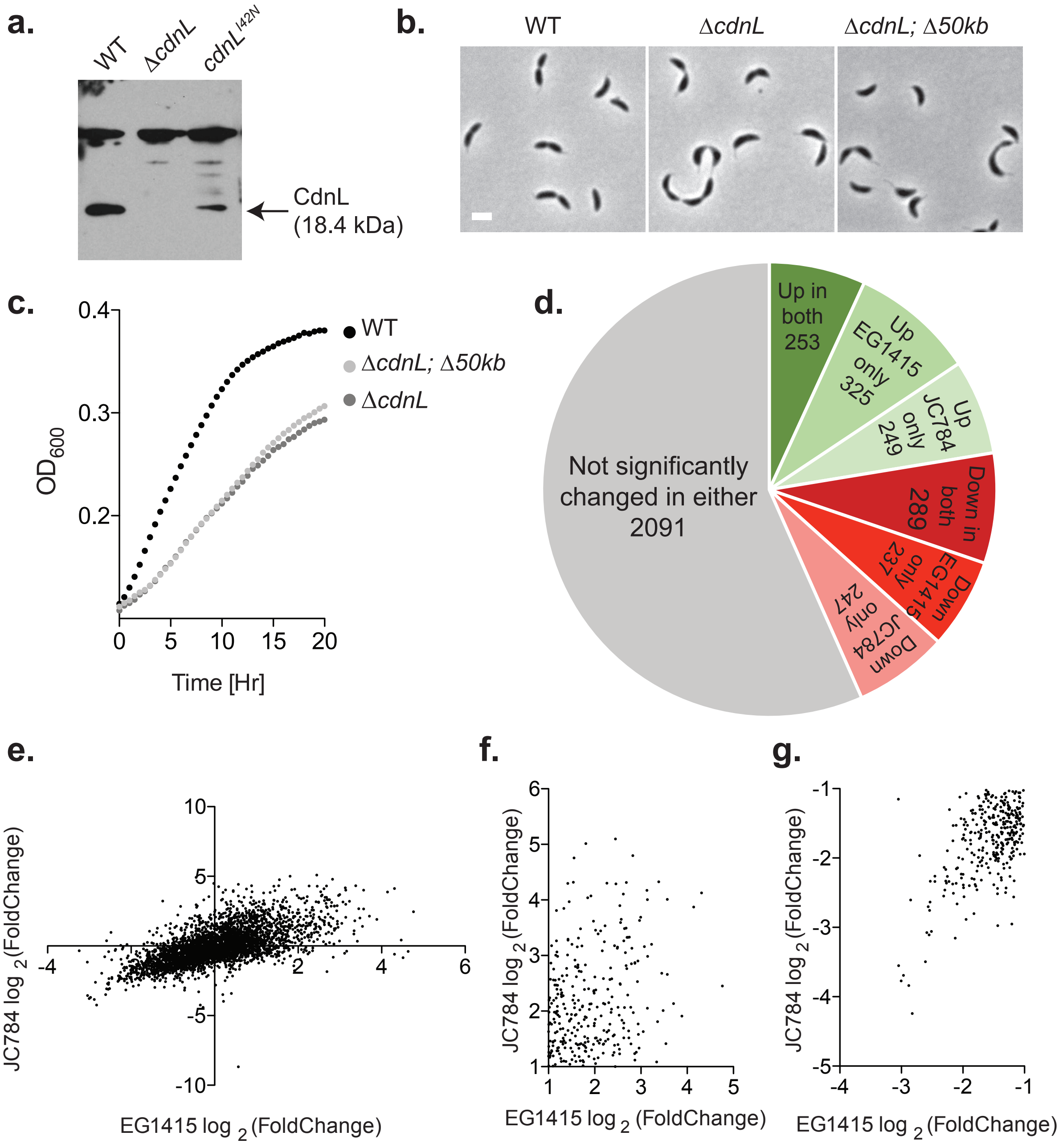


**Supplementary figure 1: RNA-Seq and microarray analysis using strains EG1415 and JC784 have significant overlap. a.** Immunoblot against whole cell lysates of the indicated strains probed with CdnL antisera. **b.** Phase contrast images and **c.** growth curves of WT

(EG865), Δ*cdnL* (EG1447) and Δ*cdnL;* Δ*50kb* (EG1415). Bar = 2 μm. **d.** Pie chart comparing RNA-Seq data using EG1415 and microarray data using JC784. A 2-fold change in transcript level was used as the cut-off for up- or downregulation. **e.** Pearson's correlation between microarray (y – axis) and RNA-Seq (x – axis) data. Pearson r = 0.6331; significant correlation with alpha = 0.05. **f & g.** Zoomed in scatter plots depicting correlation between microarray (y – axis) and RNA-Seq (x – axis) for ≥2-fold upregulated genes **(f)** and ≥2-fold down regulated **(g)** genes in both data sets.

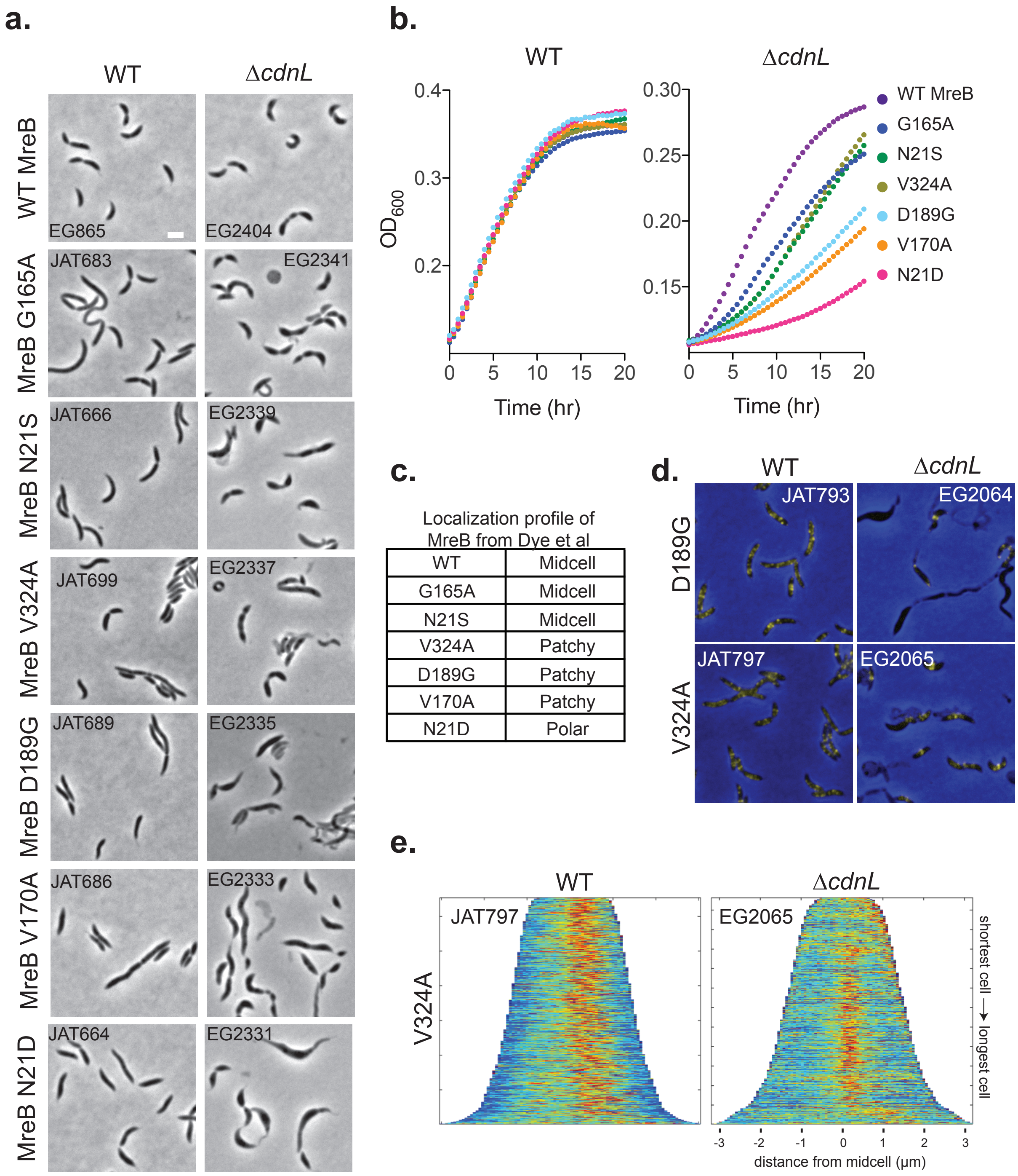


| WT | Midcell |
|---|---|
| G165A | Midcell |
| N21S | Midcell |
| V324A | Patchy |
| D189G | Patchy |
| V170A | Patchy |
| N21D | Polar |

**Supplementary figure 2: *mreB* mutants are synthetic sick with Δ*cdnL*. a.** Phase contrast images of WT and Δ*cdnL* cells expressing indicated *mreB* mutants at the *mreB* locus. Bar = 2 µm. **b.** Growth curves of strains shown in **(a)** grown in PYE. **c.** Summary of the localization of MreB mutants from Dye et al, 2011[1]. **d.** Localization of indicated MreB mutants by expressing mutant version of *venus-mreB* from *xylX* locus. **e.** Demographs of MreB mutants shown in **(d).**

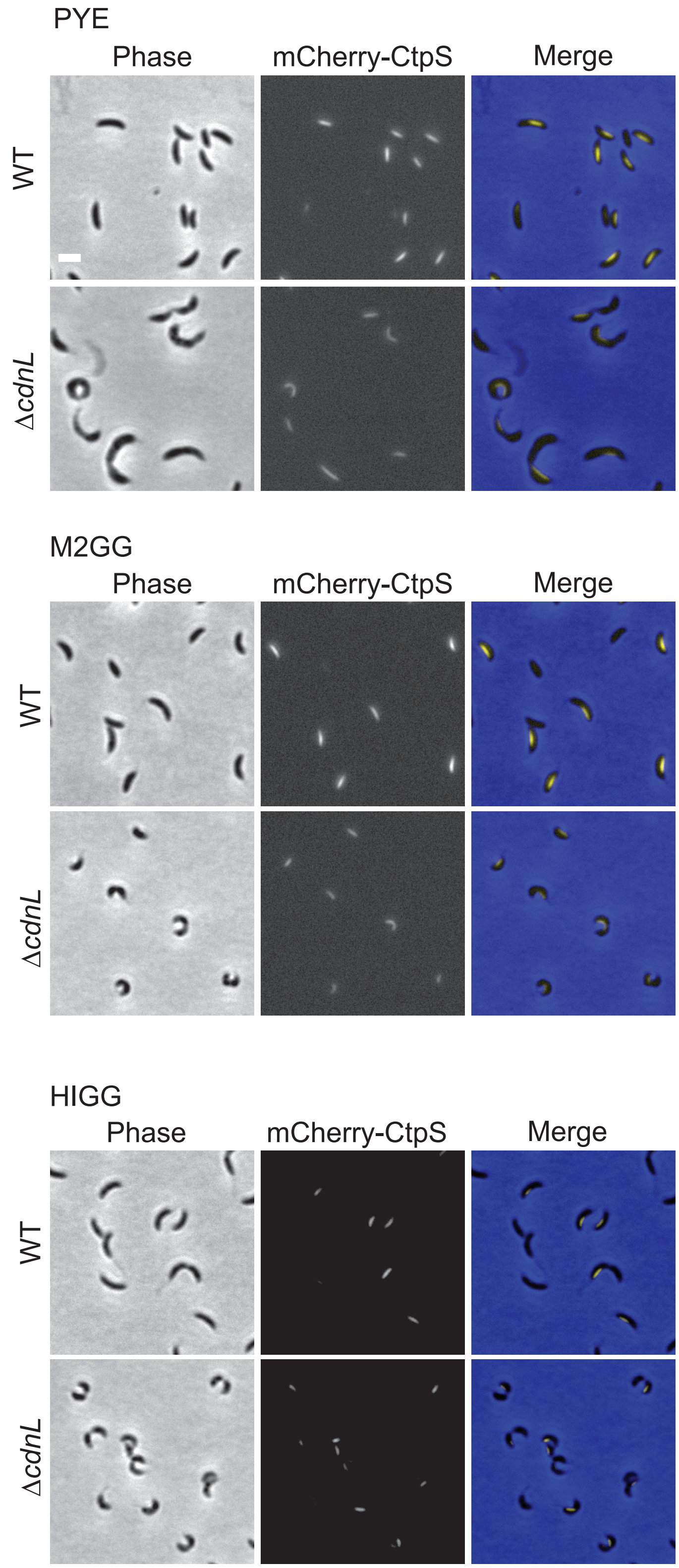
PYE
Phase
mCherry-CtpS
Merge
WT
ΔcdnL
M2GG
Phase
mCherry-CtpS
Merge
WT
ΔcdnL
HIGG
Phase
mCherry-CtpS
Merge
WT
ΔcdnL

**Supplementary figure 3. CTP synthase localization is weak in Δ*cdnL* cells compared to WT**.

Phase contrast, fluorescent and merged images of WT (EG1902) and Δ*cdnL* (EG1958) cells expressing *mCherry – ctpS* from the *xylX* locus grown in indicated media. Bar = 2 µm.

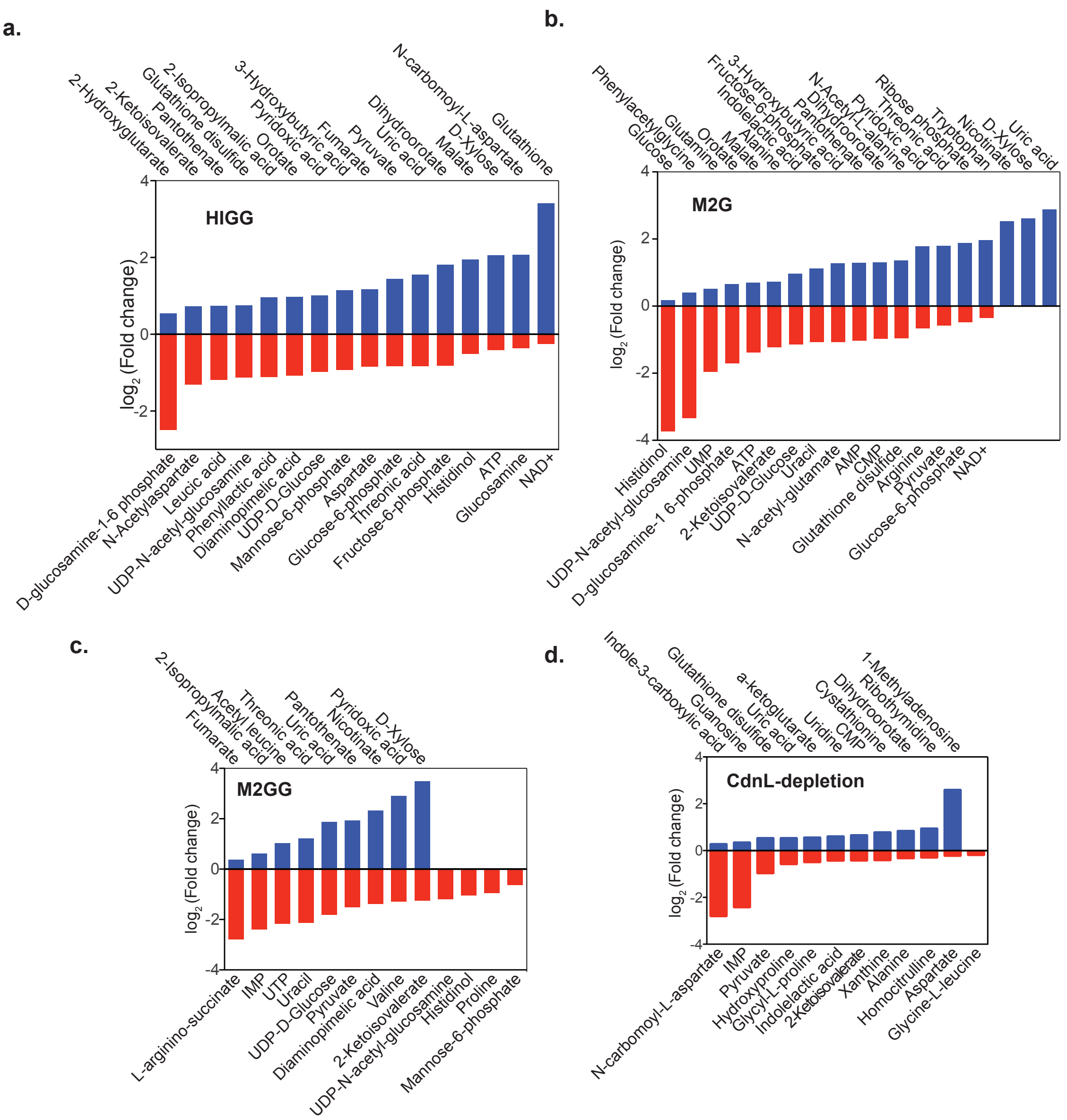


**Supplementary figure 4: Significantly altered metabolites grown in indicated media from Supplementary table 3. a-d.** WT and Δ*cdnL* cells were grown in indicated media until log-phase. **b.** Cells were grown in M2GG, washed and grown in M2G for 12 hours before extracting metabolites. **d.** Cells were grown without xylose for 8 hours to deplete CdnL before metabolite extraction. $p < 0.05$

**a.**

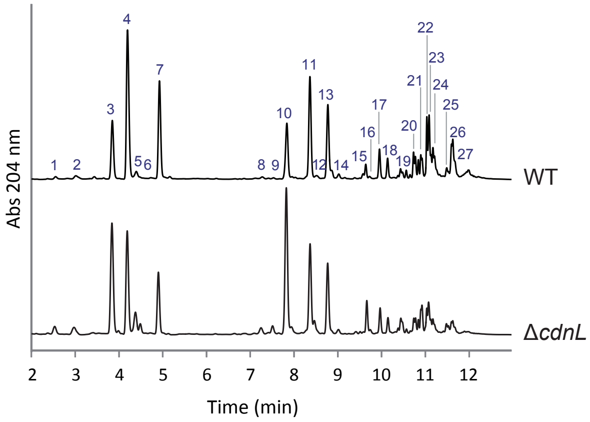


**b.**

| Parameter | WT | Δ*cdnL* |
|---|---|---|
| Monomers | 60.66 | 60.23 |
| Dimers | 26.23 | 32.22* |
| Trimers | 13.10 | 7.55* |
| Crosslink | 52.44 | 47.33 |
| Anhydro | 17.51 | 13.74* |
| Gly muropep. | 21.66 | 43.54* |
| Pentapeptides | 49.39 | 59.69* |

**c.**

| Peak | Muropeptide | Structure | WT | Δ*cdnL* |
|---|---|---|---|---|
| 1 | M3 | GlcNAc-MurNAc-L-Ala-D-Glu-mDAP | 0.60 | 1.41 * |
| 2 | M4$^{G}$ | GlcNAc-MurNAc-L-Ala-D-Glu-mDAP-Gly | 0.98 | 1.89 * |
| 3 | M5$^{G}$ | GlcNAc-MurNAc-L-Ala-D-Glu-mDAP-D-Ala-Gly | 10.69 | 20.39 * |
| 4 | M4 | GlcNAc-MurNAc-L-Ala-D-Glu-mDAP-D-Ala | 27.40 | 16.88 * |
| 5 | M2 | GlcNAc-MurNAc-L-Ala-D-Glu | 1.82 | 4.37 * |
| 6 | M44$^{G}$ | M4-M4$^{G}$ (DD-crosslink, dissaccharide lost) | 0.00 | 1.54 * |
| 7 | M5 | GlcNAc-MurNAc-L-Ala-D-Glu-mDAP-D-Ala-D-Ala | 17.22 | 9.94 * |
| 8 | D44$^{G}$ | M4-M4$^{G}$ (DD-crosslink) | 0.24 | 0.64 * |
| 9 | D43 | M4-M3 (DD-crosslink) | 0.16 | 0.75 * |
| 10 | D45$^{G}$ | M4-M5$^{G}$ (DD-crosslink) | 5.18 | 12.50 * |
| 11 | D44 | M4-M4 (DD-crosslink) | 8.80 | 7.07 * |
| 12 | M5$^{G\ Anh}$ | GlcNAc-(1-6anhydro)MurNAc-L-Ala-D-Glu-mDAP-D-Ala-Gly | 0.75 | 2.37 * |
| 13 | D45 | M4-M5 (DD-crosslink) | 6.59 | 5.88 |
| 14 | M4$^{Anh}$ | GlcNAc-(1-6anhydro)MurNAc-L-Ala-D-Glu-mDAP-D-Ala | 0.87 | 0.93 |
| 15 | T445$^{G}$ | M4-M4-M5$^{G}$ (DD-crosslink) | 0.63 | 1.35 * |
| 16 | M5$^{Anh}$ | GlcNAc-(1-6anhydro)MurNAc-L-Ala-D-Glu-mDAP-D-Ala-D-Ala | 0.33 | 0.51 * |
| 17 | T444 | M4-M4-M4 (DD-crosslink) | 1.27 | 0.98* |
| 18 | T445 | M4-M4-M5 (DD-crosslink) | 0.89 | 0.68 * |
| 19 | D45$^{G\ Anh}$ | M4-M5$^{G\ Anh}$ (DD-crosslink) | 0.92 | 1.57 * |
| 20 | D44$^{Anh}$ | M4-M4$^{Anh}$ (DD-crosslink) | 2.41 | 1.57 * |
| 21 | D45$^{Anh}$ | M4-M5$^{Anh}$ (DD-crosslink) | 1.93 | 2.25 * |
| 22 | T445$^{G\ Anh}$ | M4-M4-M5$^{G\ Anh}$ (DD-crosslink) | 1.77 | 0.73 * |
| 23 | T444$^{Anh}$ | M4-M4-M4$^{Anh}$ (DD-crosslink) | 3.04 | 1.57 * |
| 24 | T445$^{Anh}$ | M4-M4-M5$^{Anh}$ (DD-crosslink) | 1.99 | 0.76 * |
| 25 | T445$^{G\ Anh,\ Anh}$ | M4-M4$^{Anh}$-M5$^{G\ Anh}$ (DD-crosslink) | 0.50 | 0.57 |
| 26 | T444$^{Anh,\ Anh}$ | M4-M4$^{Anh}$-M4$^{Anh}$ (DD-crosslink) | 3.02 | 0.72 * |
| 27 | T445$^{Anh,\ Anh}$ | M4-M4$^{Anh}$-M5$^{Anh}$ (DD-crosslink) | 0.00 | 0.20 * |

**Supplementary figure 5: Muropeptide composition is altered in Δ*cdnL* cells. a.** Representative chromatograms of the muramidase-digested sacculi of WT and Δ*cdnL* cells. **b.** Relative molar abundance (%) of monomers, dimers, trimers, percentage of crosslinkage (proportion of crosslinked peptide side chains), muropeptides with a residue of (1-6 anhydro) N-acetyl muramic acid (Anhydro), Gly containing muropeptides (Gly muropep.), and pentapeptides. **c.** Muropeptide relative molar abundance (%). GlcNAc: *N*-Acetyl glucosamine. MurNAc: *N*-Acetyl muramic acid. Ala: Alanine. Glu: Glutamic acid. mDAP: meso-diaminopimelic acid. Gly: Glycine. Statistical analysis performed using t-test analysis. * = p < 0.05 and > 10% variation compared to WT.

**Supplementary table legends**

**Supplementary table 1**: RNA-Seq and microarray data for EG1415 and JC784, respectively. Gene (CCNA), annotation, fold change in Δ*cdnL* over wild-type, $\log_2$ of fold change, and statistical significance (1 = significant, 0 = not significant) are presented for each dataset. The 50 kb deletion in EG1415 is highlighted.

**Supplementary table 2**: DAVID[2] functional analysis of the set of genes great than or equal to 2-fold up-regulated or 2-fold down-regulated in both RNA-Seq and microarray datasets from supplementary table 1. Functional annotation clusters identified, their enrichment scores, cluster names, and the gene numbers for that cluster are indicated.

**Supplementary table 3:** Metabolomics data for EG1447 (Δ*cdnL*), EG865 (WT), EG1403 (Δ*cdnL*, *xylX::cdnL*) and EG2740 (*xylX::cfp*) grown in indicated strains.

**Supplementary table 4**: Tn-Seq data for EG1447 (ΔcdnL) and EG865 (WT). The number of unique transposon insertions in each gene in wild-type compared to Δ*cdnL* was used to determine the $\log_2$ fold-change in number of unique transposon insertions (logFC), log counts per million reads (logCPM), PValue, false-discovery rate (FDR), and negative $\log_{10}$ of the FDR for each gene in WT over Δ*cdnL*.

**Supplementary table 5:** Strains and plasmids used in this study.
